# Design of artificial molecular motor inheriting directionality and scalability

**DOI:** 10.1101/2023.07.19.549658

**Authors:** Kenta I. Ito, Yusuke Sato, Shoichi Toyabe

## Abstract

Realizing artificial molecular motors with autonomous functionality and high performance is a major challenge in biophysics. Such motors not only provide new perspectives in biotechnology but also offer a novel approach for the bottom-up elucidation of biological molecular motors. Directionality and scalability are critical factors for practical applications. However, the simultaneous realization of both remains challenging. In this study, we propose a novel design for a rotary motor that can be fabricated using a currently available technology, DNA origami, and validate its functionality through simulations with practical parameters. We demonstrate that the motor rotates unidirectionally and processively in the direction defined by structural asymmetry, which induces kinetic asymmetry in motor motion. The motor also exhibits scalability, such that increasing the number of connections between the motor and stator allows for a larger speed, run length, and stall force.

**Statement of Significance:** Biological molecular motors are the most sophisticated nanomachines found in nature. Despite a long history of research, we have yet to understand how they work in detail. A simple and promising approach involves the building of artificial molecular motors. The process of achieving these goals helps us understand how the structures and mechanisms of the biological motors have been optimized through the evolution of their functions. Some artificial motors have already been achieved. However, each realization implements only a part of the characteristics of biological motors. In this work, we propose a novel motor mechanism that simultaneously implements two essential characteristics: scalability and directionality. These results will help us improve the biophysics of molecular motors and contribute to developing high-performance artificial molecular motors with possible industrial applications.

## Introduction

Artificial molecular motors with autonomous functionalities are expected to open up novel dimensions in biophysics^1,2^. They may serve as nano-sized chemically-fueled actuators for, for example, artificial muscle. Attempts to realize artificial molecular motors are expected to improve our understanding of chemo-mechanical coupling at the nanoscale and advance biophysics. A significant trend in fabricating artificial molecular motors is to exploit DNA nanotechnology with the help of nuclease enzymes. The enzyme cleaves fuel DNA or RNA strands to liberate free energy required for driving motors^1^.

A molecular motor must implement three ingredients for unidirectional motion: driving force, chemo-mechanical coupling, and kinetic asymmetry. The driving force is the free energy liberated by, for example, the cleavage of fuel molecules. The chemo-mechanical cycle of the motor couples the free energy consumption and unidirectional motion. Since the physical laws governing the motor motion are spatially symmetric, the translation direction is not defined without kinetic asymmetry, that is, the bias to prefer transition in a specific direction^3^. An additional ingredient necessary for practical applications is the scalability, or the ability of multiple units to work in parallel so that the device amplifies the velocity and force. A prominent example is skeletal muscle^4^, which consists of thousands of myosin II motors and realizes macroscopic contraction amounting to the length of ∼ cm and the force of ∼ N.

Generally, a translational molecular motor performs a unidirectional motion by dissociating at its rear side from the stator and binding at its front side to the stator. Biological molecular motors such as kinesin and myosin V realize such kinetic asymmetry even though their rear and front legs (often called heads) are identical^5,6^. They exploit a conformational change of the legs to create an asymmetric leg-rail interaction. However, the synthetic motors created so far have exploited simpler mechanisms to achieve a forward-to-backward bias. The two primary biasing mechanisms are the burnt-bridge Brownian ratchet (BBR) and bi-pedal motors.

The BBR motor irreversibly disrupts rail molecules as fuel at the rear so that the motor inevitably moves forward by blocking the backward motion^7–12^. Thus, the motor creates a fuel gradient in the rail and translates by self-avoiding the Brownian motion. An outstanding feature of the BBR motors is their high scalability. Their simple chemo-mechanical mechanism allows one to increase the number of connections between the motor and stator working in parallel to improve speed and force. However, BBR motors have an inherent limitation in directionality owing to the lack of internal mechanisms to create kinetic asymmetry; in principle, they translate in the direction determined by the first step, which is randomly chosen by fluctuations. Therefore, the ensemble-averaged speed vanishes unless the initial state is biased, as realized previously^8^. In addition, because they cleave the fuel molecules that serve as a part of the stator, a rotational mechanism in which the rotor iteratively passes through each stator location is not feasible.

The bi-pedal motors are able to walk on a track by a hand-over-hand mechanism in the direction defined by structural asymmetry, which generates kinetic asymmetry^13–15^. The motion is processive because the fuel is supplied from the solution, and the motor does not disrupt the stator. However, a scalable extension necessary for practical use is not straightforward.

This work proposes a novel motor mechanism that simultaneously inherits directionality and scalability and implements the mechanism in a rotary motor. The rotor moves autonomously and processively in a cylindrical stator in a direction determined by the kinetic asymmetry induced by the stator’s structural asymmetry. The driving force is obtained by nicking the fuel DNA strands connecting the cylinder and rotor by nicking enzyme dispersed in the solution. Rotor’s motion combines rotations around the rotor axis and orbital revolutions along the inner side of the stator. The stator implements fuel replacement, which allows processive motion of the rotor. The structure and chemo-mechanical mechanism are designed such that the motor could be fabricated using DNA origami technology^16,17^. To validate the concept, we conducted extensive numerical simulations with practical physical parameters.

## Brief methods

### Motor design

The motor consists of a rotor shaft with a square-pillar shape and a stator with a hexagonal-cylinder shape (Fig. 1). We designed the structure using caDNAno, software for designing DNA-origami structures, to ensure that the structure is fabricable by DNA nanotechnology^18^ (Fig. 1a). Geometrical parameters obtained by caDNAno were used for the simulations (Table 1).

**Table 1.** Geometrical and reaction parameters.

| Description | Label | Value | Unit |
| --- | --- | --- | --- |
| Inner diameter of stator | - | 31 | nm |
| Outer diameter of rotor | - | 7.5 | nm |
| Height of stator and rotor | - | 70 | nm |
| Number of S and R strands in a row | $N$ | 1 ~ 15 | |
| Equilibrium length of S-F-R connection | $L_{\text{eq}}$ | 8 | nm |
| Spring constant of S-F-R connection | $k$ | 24.7 | $k_B T / \text{nm}^2$ |
| Equilibrium angle of S | $\phi_{\text{eq}}$ | 45 | degree |
| Angular spring constant of S (angular) | $k_{\theta}$ | $64/\pi^2$ | $k_B T$ |
| Debye length | $\lambda_D$ | 1.25 | nm |
| Cut-off length | $r_{\text{cutoff}}$ | 12 | nm |
| $\Delta G$ of S-F hybridization in solution | $\Delta G_{\text{chem}}^{\text{S}}$ | -1 | $k_B T$ |
| $\Delta G$ of R-F hybridization in solution | $\Delta G_{\text{chem}}^{\text{R}}$ | 2 | $k_B T$ |
| $\Delta G$ of R-decomposed product hybridization | - | 6 | $k_B T$ |
| Concentration of F | - | 3 | $\mu\text{M}$ |
| Binding rate of two tethered strands within $r_{\text{cutoff}}$ | $k_{\text{bind}}^0$ | 500 | $\text{s}^{-1}$ |
| Binding rate of free F in solution to R and S | $k_{\text{on}}$ | 1 | $\text{s}^{-1}$ |
| Enzymatic cleavage rate | $k_{\text{cut}}$ | 0.0015–1.5 | $\text{s}^{-1}$ |

**Figure 1.**
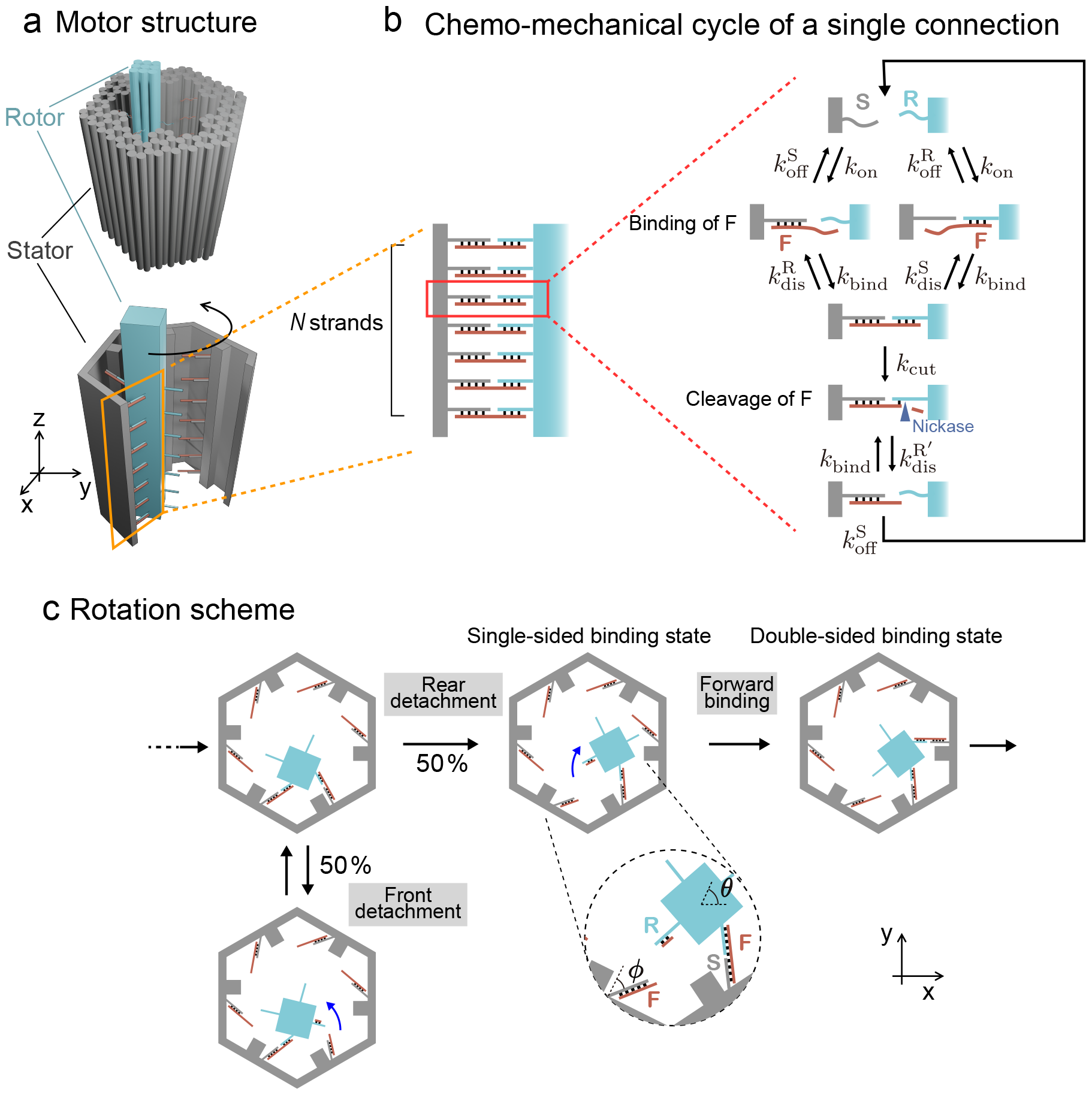
The design of the motor structure and reaction scheme. **a**, Schematic of structure designed by caDNAno (top) and the coarse-grained structure used for the simulation (bottom). A side wall of the stator is removed so that the internal structure is visible. **b**, Chemo-mechanical cycle of a single connection. Only a single set of S and R is shown for simplicity, although multiple sets exist. **c**, Rotation scheme in a cross view. Although there are *N* strands in each row, only a single set of strands is shown at each side for simplicity.

*N* single DNA strands are arrangd in rows on each side of the rotor and stator (Fig. 1b). The motor alternatively iterates a state with connections on a single side and a state with connections on double sides (Fig. 1c). We do not consider states with connections on triple sides simultaneously since they can be prevented by carefully designing the structures in the experiments.

The strands on the rotor and stator are denoted R and S, respectively. Steric hindrances on the stator bias the stable position of angle *ϕ* of S by electrostatic repulsion and introduce structural asymmetry.

The fuel DNA strands F dispersed in the solution serve as a linker connecting the rotor and stator. The sequence of F is designed such that each end of F hybridizes with R and S. An F-R binding is unstable, whereas an F-S binding is stable. Hence, most R is single-stranded, while most S is bound by F (Fig. 1c). Due to the instability of the F-R binding, a single S-F-R connection is unstable. However, multiple S-F-R connections are stabilized; even if a connection between two bodies is broken, the other connections keep the two bodies close, which increases the re-connection rate of the broken connection. Thus, the stability increases.

A nicking enzyme dispersed in the solution cleaves the F hybridized to R. The nicking site is located in the region where F and R hybridize (Fig. 1b). Since F-R binding is unstable, cleavage is basically limited to the F strands in the S-F-R connections. Consequently, the cleavage reactions of F depend on the rotor-stator conformation, where the F strands connect the rotor and stator, and the conformation is changed by the cleavage reactions. Thus, the tuned binding stability between F, S, and R implements chemo-mechanical coupling. The cleavage of F provides a free energy change necessary for driving a unidirectional motion. The cleaved products of F, remaining on R and S, readily dissociate. Then, an intact F strand dispersed in the solution binds to S.

The mechanism will be elaborated in the results section.

### Simulation overview

We developed a stochastic simulation model based on the Gillespie algorithm^19^. The system is complicated in that chemical and mechanical degrees of freedom interact. However, the thermal relaxation time of the mechanical degrees of freedom, including the motor position and angles of S and R, is typically much faster than the time scales of chemical dynamics, including hybridization, dissociation, and fuel cleavage. Therefore, we reasonably assumed that the mechanical degrees of freedom are always equilibrated.

We simulated stochastic chemical transitions among distinct chemical states, in which the mechanical degrees of freedom are locally equilibrated, by the Gillespie algorithm. The Gillespie algorithm reproduces the exact dynamics of stochastic Markovian systems with given reaction rates. The reaction rates of the chemical transitions are determined based on the equilibrium distribution of the mechanical degrees of freedom described by the coordinates *x* and *y* and the rotational angle *θ* of the motor (Eq. 6) and the chemical properties of the system under the constraint of the local detailed balance.

The mechanical degrees of freedom are determined as the stable distribution for each chemical state. Thus, we did not explicitly calculate the dynamics of the mechanical degrees of freedom.

More details have been provided in the Methods section.

## Results

### Directionality

We explored the conditions for the motor to translate unidirectionally, explicitly focusing on the dependence on the cleavage rate *k*_cut_. We observed three distinct operational modes depending on *k*_cut_; self-directed unidirectional motion, Brownian motion, and BBR motion (Fig. 2a–d). Here, we define the unidirectional motion as the motion with the finite ensemble-averaged velocity larger than the standard deviation of the velocity. As mentioned, the BBR shows a ballistic motion on a short time scale; however, the direction is chosen randomly. Thus, the ensemble-averaged velocity vanishes for the naive BBR mechanism.

**Figure 2.**
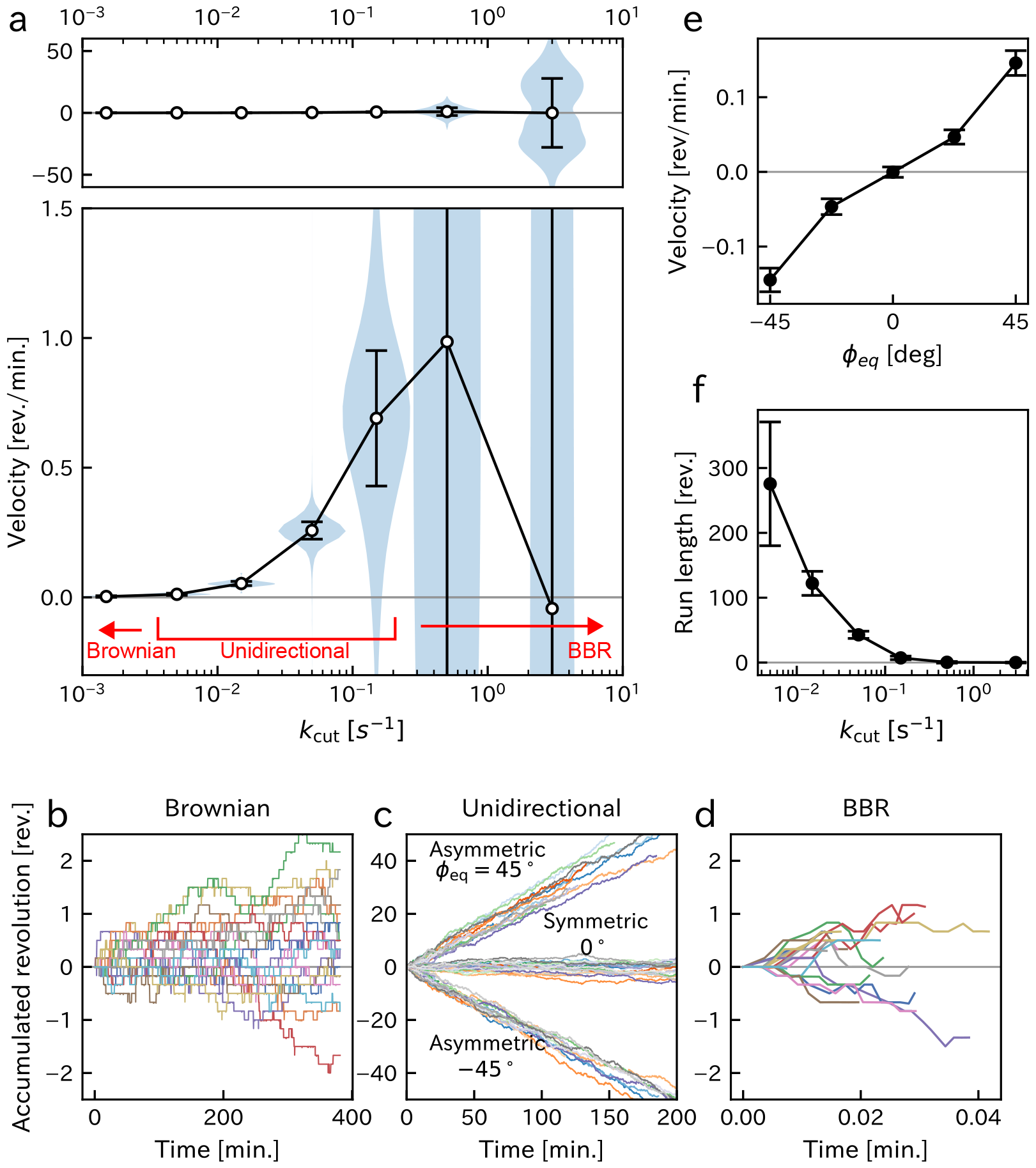
Conditions for unidirectional motion. **a**, Mean velocity with the violin plot indicating the frequency by width. The bottom panel shows the magnification. The numbers of runs are 4986, 999, 999, 908, 70, 70, 70, respectively, from right to left. **b-d**, Simulated trajectories of the revolutions (20 runs in each group) with *k*_cut_ = 5 × 10^−5^ s^−1^ (b), 0.05 s^−1^ (c), and 1.5 s^−1^ (d). Different colors correspond to independent simulations. *ϕ*_eq_ = 45° except for **c**. In **c**, *ϕ*_eq_ = 45° (top), 0° (middle), and − 45° (bottom) are compared. **e**, Average velocity vs *ϕ*_eq_. *k*_cut_ = 0.05 s^−1^. The number of runs is 50 for each point. **f**, Run length vs *k*_cut_. *N* = 6. Error bars in **a, e**, and **f** represent standard deviations.

### Self-directed unidirectional motion

We observed a unidirectional motion by tuning *k*_cut_ to intermediate values significantly less than the binding rate of F to R and S, *k*_on_ = 1 s^−1^ (Fig. 2a, c). Such unidirectional motion was not observed without structural asymmetry (Fig. 2c, e) or fuel consumption with a vanishing *k*_cut_ (Fig. 2a, b). The rotation direction was inverted with opposite structural asymmetry (Fig. 2c, e). These results validate that the motor is driven by the free energy consumption in the direction defined by the kinetic asymmetry induced by the structural asymmetry. Note that the motor revolves in the stator while rotating on its own axis.

The unidirectional motion suggests a biasing mechanism that favors the dissociation of the rear side in the double-sided binding state, binding at the front side in the single-sided binding state, or both. These possibilities will be examined in the following.

Under the present conditions, with intermediate *k*_cut_ vlues, the connections by F are mainly broken by enzymatic cleavage rather than by dehybridization. The fuel cleavage rates on the front and rear sides are the same in the double-sided binding state. Since *k*_on_ *> k*_cut_, the cleavage of all connections on a side requires an extended duration, which is sufficient for each connection to repeat multiple cycles of fuel binding, cleavage, and replacement before dissociation. Therefore, the number of connections on each side fluctuates in a random-walk manner. In the present setup, the dissociation rate of each connection is almost the same, independent of the side to which it belongs. Consequently, the front and rear sides dissociate stochastically at similar rates. That is, there is no forward-to-backward bias in the dissociation process unless there is a bias in the initial distribution of connections. Such a biased distribution is not expected under the present conditions with relatively small *k*_cut_ values.

In contrast, in the single-sided binding state, steric hindrance tilts the motor forward and makes forward binding more feasible than backward binding. The speed increased with |*ϕ*_eq_| because of the enhanced binding bias as far as |*ϕ*_eq_| is relatively small. When |*ϕ*_eq_| is too large, the binding of the rotor and stator via two sides becomes unstable, resulting in the fast dissociation of the motor (data not shown). When *ϕ*_eq_ is large in negative, the distance for forward binding increases. When *ϕ*_eq_ is large in positive, the rotor feels a strong repulsion from the stator cylinder.

Thus, only binding is biased in the forward direction. Hence, on average, the motor performs a forward step every approximately two chemo-mechanical cycles. All connections on a side may dissociate even in the single-sided binding state, resulting in the decomposition of the motor-stator complex.

The speed increased with *k*_cut_ in the unidirectional-motion regime (Fig. 2a) since dissociation is the rate-limiting process. The average run length, evaluated as the average traveling distance until both sides dissociate from the stator, decreases with *k*_cut_ due to the increase in the dissociation rate in the single-sided binding state (Fig. 2f). This indicates a trade-off between speed and processivity.

### Brownian motion

At negligible *k*_cut_, the motor exhibited Brownian motion with a vanishing mean speed (Fig. 2a, b). As the driving force is absent at the vanishing *k*_cut_, unidirectional motion is forbidden by the microscopic reversibility^20^. In other words, under these conditions, with a negligible fuel cleavage rate, dissociation is driven by the dehybridization of F from S or R. Therefore, binding to the forward side and dissociation of the forward side are opposite reactions. Although forward binding is still more favorable than backward binding because of structural asymmetry, dissociation has the same bias. This cancels out the binding bias.

### BBR motion

When we increased *k*_cut_ to the values comparable to the fuel binding rate *k*_on_ = 1 s^−1^, a unidirectional motion was still observed (Fig. 2d). With such a large *k*_cut_, fuel cleavage proceeds faster than fuel replacement, keeping the fraction of S possessing F small. In the single-sided binding state, the backward step is prevented due to the lack of available intact F strands on S; the binding is biased in the forward direction, resulting in a unidirectional motion.

However, the rotational direction was random and varied from run to run, as indicated by in the symmetrical two-peak frequency distribution (Fig. 2a). The ensemble-average speed vanished despite structural asymmetry, and the variance was significant (Fig. 2a). Furthermore, the rotation was not processive; the motor readily dissociated from the stator after a few revolutions (Fig. 2d, f). Fast fuel cleavage reduces the amount of intact fuel strands on the stator and prevents the motor from reconnecting to the stator. Therefore, the rotor easily dissociates from the stator. A processive revolution is not achieved unless the cleavage is slower than the fuel replacement. These are the distinct characteristics of the BBR-type mechanism.

### Scalability

We evaluated the scalability of the unidirectional motion (*k*_cut_ = 005*s*^−1^) in terms of the *N* dependence of the velocity, run time, and run length. The run time and length quantify the motor processivity and are defined as the time duration and traveling distance, respectively, before both sides dissociate from the stator.

We found that the velocity decreased with *N* (Fig. 3a), whereas the run time and length increased with *N* (Fig. 3b, c). As shown, the velocity increases with *k*_cut_ in the unidirectional motion regime (Fig. 2a). However, the motor rapidly dissociated at large *k*_cut_, which decreased the run length (Fig. 2f). This trade-off between the velocity and run length (Fig. 3e) practically limits the motor performance. Nonetheless, by increasing *N*, we can suppress dissociation and raise the threshold for *k*_cut_ to further increase the velocity. Thus, the motor inherits scalability, such that an increase in *N* allows a larger velocity and run length.

**Figure 3.**
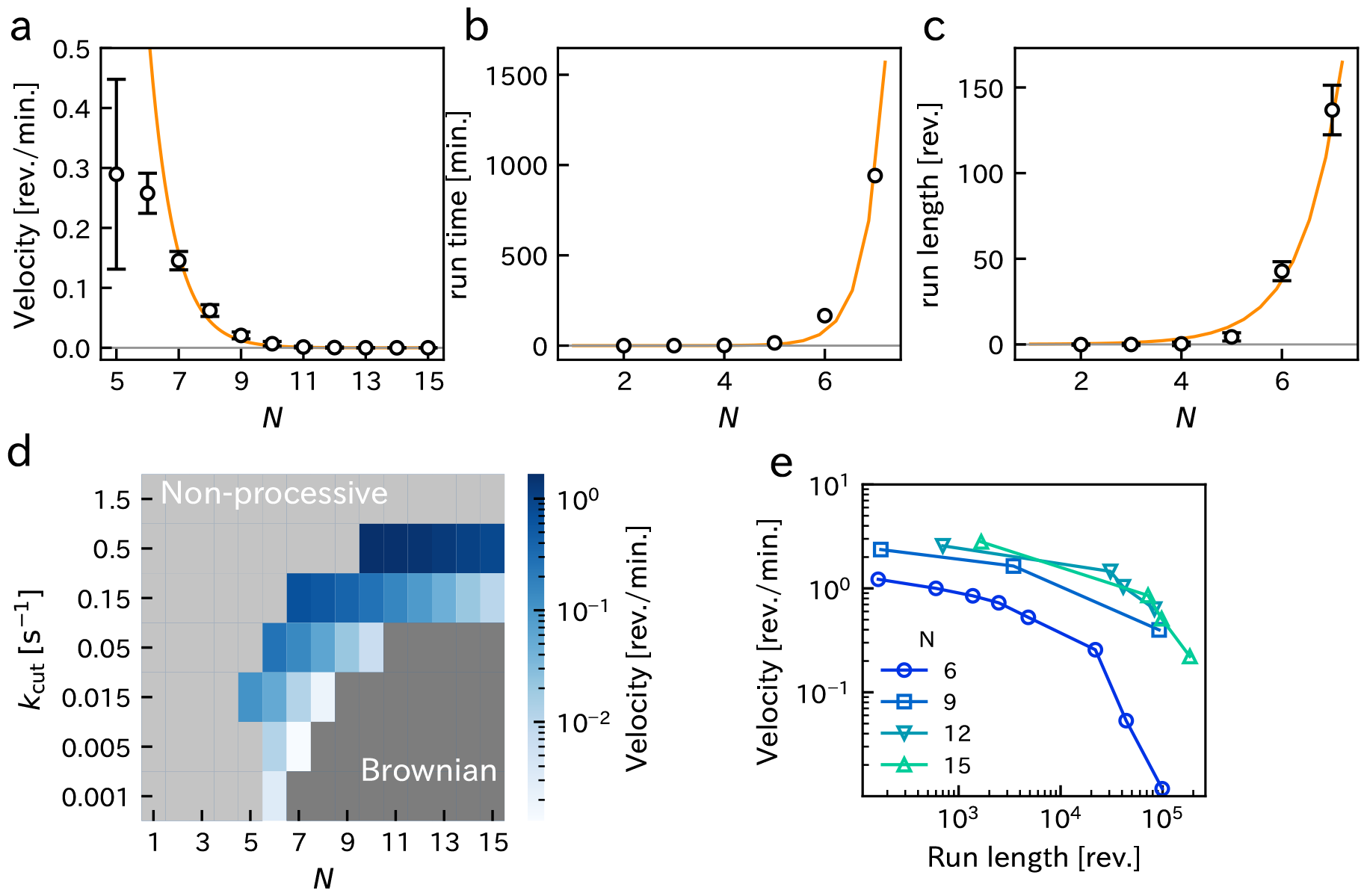
Dependence of velocity and run length on *N*. **a**, Velocities. Velocities for *N* ≤ 4 are not shown because the errors were too large. **b**, Run time. The dissociation is too fast with *N* = 1, and dissociation did not occurs with *N* ≥ 8. **c**, Run length. The solid lines in **a** – **c** are obtained by the global fitting of *aN*(1 + *K*)^−*N*^, *b/N*^2^*/*(1 + *K*)^−2*N*^, and *c/N/*(1 + *K*)^−*N*^, respectively, with *a* = 402, *b* = 0.00015, *c* = 0.050, and *K* = 3.1 (see Eq. 2). **d** Velocity as the function of *k*_cut_ and *N*. The light gray region corresponds to a non-processive motion (the run length is less than 10 revolutions). The dark gray region corresponds to Brownian motion without unidirectional motion. **e**, Velocities vs run length. Points in the same curve have different *k*_cut_ values. Error bars in **a** – **c** indicate the standard deviations. *k*_cut_ = 0.05 s^−1^ for **a** – **c**. The number of runs is 70 for each point.

The rate-limiting process of the chemo-mechanical cycle is the dissociation of a side, which requires the cleavages of all connections in the side. Therefore, it is reasonable that the increase in *N* slows down the motor while enhancing processivity. We derive the explicit *N* dependence of the velocity, run time, and run length based on a simple estimation of the dissociation rate of a single side. For simplicity, the binding dynamics of each connection are assumed equilibrated. Let *p* be the probability that a pair of R and S on a side are connected by F. Let *k*_+_ and *k*_−_ be the binding and dissociation rates of a single connection, respectively. These phenomenological parameters are complex functions of the elementary reaction parameters *k*_on_, *k*_off_, *k*_bind_, *k*_dis_, and *k*_cut_ on multiple sides of the rotor and stator. Hence, it is difficult to derive the values of *k*_±_ from other parameters. We obtain *p* = *k*_+_*/*(*k*_+_ + *k*_−_) = *K/*(1 + *K*) by solving the detailed balance *pk*_−_ = (1 − *p*)*k*_+_, where *K* ≡ *k*_+_*/k*_−_. The probability that there are *n* (0 ≤ *n* ≤ *N*) connections in the side obeys the binomial distribution, such that

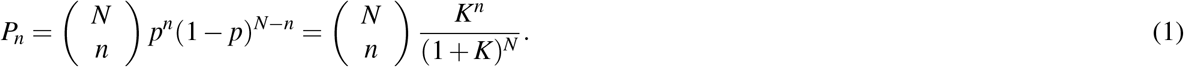

The dissociation rate of a side is calculated as

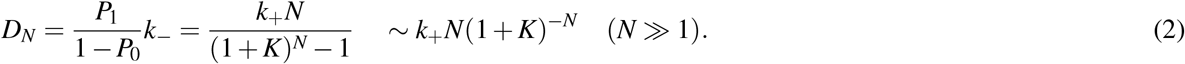

Here, 1− *P*_0_ is a normalization factor and corresponds to the probability that the side is bound.

A coarse scaling discussion implies that the velocity is proportional to the *D*_*N*_, since the side dissociation is the rate-limiting step under the present conditions. This may explain the exponential decrease in velocity with increasing *N* for large *N* (Fig. 3a). In contrast, the run time scales as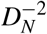, since both sides need to dissociate. The run length is calculated as the steady velocity times the run time and scales as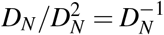. We successfully fit the velocity, run length, and run time using these scaling relations at large *N* (Fig. 3a–c).

### Stall force

Next, we evaluated the stall force of unidirectional motion (Fig. 4a, b). The stall force is the magnitude of the hindering external force necessary for stalling the motor motion and quantifies the maximum force the motor can exert. The motor motion combines rotations around the rotor axis and orbital revolutions along the inner side of the stator. Hence, we evaluated the stall force rather than stall torque.

**Figure 4.**
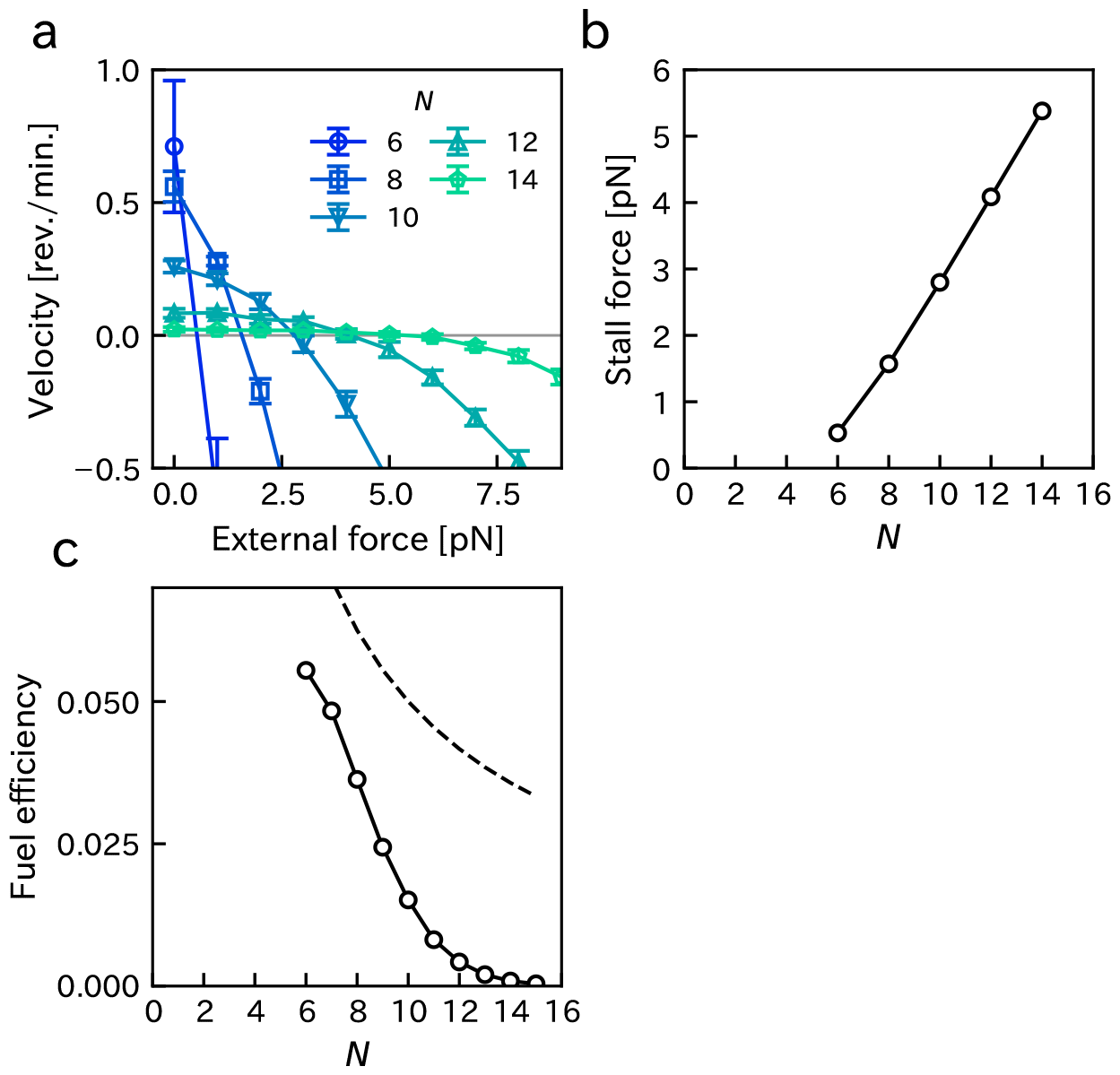
Stall force and fuel efficiency. *k*_cut_ = 0.15 s^−1^. **a**, Velocity under different magnitudes of the external force. The stall force is evaluated as the force where the velocity vanishes. Error bar indicates the standard deviation. **b**, Magnitudes of stall force for various *N*. **c**, Fuel efficiency *η* (symbol and solid line). Dashed line corresponds to 0.5*/N*, which corresponds to a situation that *N* fuel strands are consumed on average per cycle. The number of runs is 20 for each point.

We induced an external hindering force on the rotor in the direction perpendicular to the rotor revolution. As the magnitude of the applied force increases, the velocity decreases, vanishes, and then reverses (Fig. 4a). The stall force, evaluated as the external force at which the speed vanished, increased with *N* (Fig. 4b). The unidirectional motion of the motor originates from the connection’s tilt owing to steric hindrance (Fig. 2e). Therefore, reducing the tilt caused by the external hindering force is expected to slow the motor. As the hindering force is distributed over the connections, an increase in *N* suppresses the tilt reduction and increases the stall force.

We obtained a stall force of several pN (Fig. 4b), which is comparable to the stall force of an artificial bi-pedal DNA motor^14^, kinesin (5–7pN)^21^, and myosin V (2–3pN)^22^. The stall torque, which is roughly estimated to be several tens of pNnm by using 10 nm as the rotational radius, exceeds the stall torque of F_1_-ATPase (approximately 40 pNnm)^23^. The stall torque of the bacterial flagellar motor (approximately 1000 pNnm)^24^ is much larger because it is strongly driven by proton motive force. A large stall force of ∼150 pN was reported for a BBR-type motor with *>* 1000 connections^25^. However, these BBR motors do not achieve self-directed unidirectional motion.

### Fuel efficiency

Finally, we evaluated the efficiency of unidirectional motion in terms of fuel consumption. Fuel efficiency *η* is defined as the mean number of net forward steps divided by the mean number of consumed fuel molecules during a certain period (Fig. 4c).

We found that *η <* 0.5 and that *η* decreases with *N*. The decrease in *η* with *N* may be explained as follows. The cleaved fuels may be replaced by intact fuel strands before all connections on the side are cleaved. Such “resurrection” of the fuel leads to wasteful fuel consumption, while the efficiency decreases. As *N* increases, the time required for all connections on the side to be cleaved increases; therefore, futile fuel consumption increases. Since the motor steps every two cycles on average (Fig. 1c), *η* = 0.5*/N* if *N* fuel strands are consumed on average per cycle, which is indicated as a dashed line for a reference in Fig. 4c. The scaling of the efficiency and other parameters with *N* is one of the main topics in the field^26^. A study on BBR-type motors demonstrated an increase in efficiency with *N*^27^. Future studies should integrate thse mechanisms into the current system.

## Discussion

This study proposes a novel design for an artificial molecular motor with controlled directionality and scalability. Unlike BBR motors, this motor implements fuel replacements, which enables rotational motion. The structure is designed such that it is fabricable using the available technology. The motor exhibited processive unidirectional motion in the direction determined by kinetic asymmetry induced by structural asymmetry. The simplicity of this mechanism makes the motor scalable. Increasing the maximum number of connections, *N*, allows for larger velocity, run length, and stall force (Fig. 3a-c). Thus, this design has the advantages of both track-walking bi-pedal motors and multivalent BBR motors.

However, the velocity was limited to a relatively small value of ∼1 rev./min. (Fig. 4a). The fuel efficiency *η* was also limited to small values (Fig. 4c). This highlights the sophisticated chemo-mechanical coupling mechanism of biological molecular motors. To enhance motor performance, it is critical to bias also the dissociation process so that the rear side dissociates more readily compared to the front side. Since the front and rear sides are identical, we need an additional mechanism to implement such bias. Biological molecular motors such as kinesin and myosin V utilize conformational changes to achieve a significant bias for both binding and dissociation^5,6^. The possibility of implementing conformational changes in synthetic motors has been discussed^28^. In contrast, an artificial bi-pedal motor achieved bias by using the asymmetric response of double-helix denaturing in the direction of the mechanical load^13,29^. However, it is not straightforward to implement the scalability and these mechanisms simultaneously. A simple solution to this problem is to control the accessibility of the enzyme on the front and rear sides. In the present work, we assumed that the enzyme is suspended in solution and accesses the motor via diffusion. It has been established to precisely fix proteins on the targeted location of DNA origami structure^30^. Therefore, it may be possible to cleave the connections on the rear side favorably by carefully placing the enzyme on the stator since the S-F-R connections on the front and rear sides have different equilibrium angular positions (Fig. 1b).

Experimental validation of the design will be a future challenge. Rotary structures have already been fabricated using DNA origami; however, autonomous functionality has not been implemented^31,32^. One of the benefits of the rotary mechanism is that precision observation is more accessible than that of the linear mechanism since sub-micron probes attached to the rotor can magnify the rotational motion. Note that the probes do not amplify the linear motion. Issues to be solved may include the accessibility of the fuel DNA strand and enzyme to the stator cylinder and the efficient emission of the cleaved products from the cylinder. The proposed mechanism is not limited to rotary motors. Applications to other geometries, such as linear motors, are straightforward. An intriguing development includes the accumulation of many motors to build a macroscopic actuator, such as a skeletal muscle.

## Methods

The Gillespie algorithm^19^ simulates the stochastic transitions between discrete states and provides exact sampling. In the following, we define the states and then the potentials and reaction rates necessary for defining the transition rates among the states.

### States

The chemical states are defined based on the combinations of the binding states of all R and S. Each R (S) is in one of the five possible states: single-stranded, bound to free F, bound to the shorter cleaved product of F, connected to S (R) via F, or connected to S (R) via the longer cleaved product of F.

### Motor coordinates

We assumed that the rotor motion is limited to the lateral space and does not move along the z-axis (Fig. 1). Thus, the rotor geometry is specified by the rotor centroid (*x, y*) and rotational angle *θ*.

### Potential energy

We assumed that the mechanical dynamics of (*x, y, θ*) and *θ* are much faster than the chemical reaction dynamics and always equilibrated in each chemical state. Therefore, we use the equilibrium distribution of (*x, y, θ*) given by the Boltzmann distribution with a potential including the rotor-stator electrostatic repulsion, elastic extension of the double-stranded DNA, and steric hindrance for S.

DNA nanostructures are usually built in the presence of cations to suppress electrostatic repulsion among the negatively charged DNA strands. In the simulation, we used the Debye length of *λ*_D_ = 1.25 nm assuming the presence of 20 mM MgCl_2_. Although a base pair contains two charged phosphate groups, we assumed that only a single side of the phosphate group exposed at the surfaces of DNA origami is involved in electrostatic repulsion because *λ*_D_ is smaller than the diameter of the double-stranded DNA. Furthermore, we assumed that interaction among phosphate groups at the rotor and stator surfaces is limited to those within a distance smaller than *λ*_D_ and the interaction between only the nearest double strands is effective due to the Debye shielding. When two double-stranded DNA are adjacent, the phosphate group of either strand has at most 7.5 phosphate groups in the other strand within *λ*_D_. Thus, the electrostatic potential between the rotor at *x, y* and the stator leads to

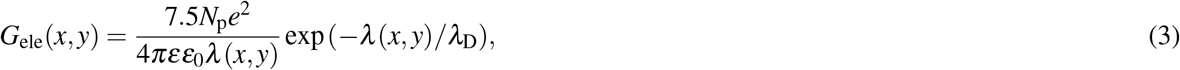

where *λ* (*x, y*) is the shortest distance between the rotor and stator cylinder under the approximation that the rotor and stator have cylindrical shapes and is calculated as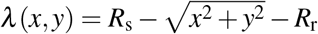. Here, *R*_s_ and *R*_r_ are the radius of the stator and rotor cylinders, respectively. *e* is the elementary charge, *ε*_0_ is the permittivity of vacuum, and *ε* is the relative permittivity of water. *N*_p_ = 210 is the number of phosphate groups in a row obtained by dividing the rotor or stator height (70 nm) by the length of a single base pair (0.33 nm).

We calculated the elastic energy of each S-F-R connection as an enthalpic linear spring with a spring constant of *k* = *ES/L*_eq_. Here, *E* is the Young’s modulus *E, S* is the cross-sectional area *S*, and *L*_eq_ = 8 nm is the equilibrium length of the double-stranded connection S-F-R. Since *L*_eq_ is much shorter than the persistence length of a double strand DNA strand (∼ 50 nm), the elasticity of the connection is reasonably assumed to be enthalpic. The typical DNA length here is 8 nm; and thus, the total elastic energy is calculated as

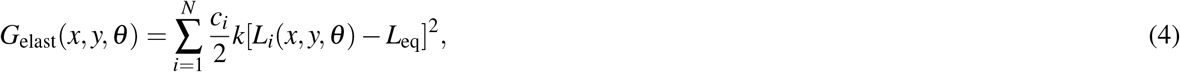

where *L*_*i*_ is the length of the *i*-th connection. *c*_*i*_ = 1 if *i*-th S and R are connected by F and 0 otherwise.

The electrostatic repulsion by the steric hindrance is modeled using a harmonic potential. Let *ϕ*_*i*_ be the angle of the *i*-th S strand with the normal direction of the cylinder’s inner surface as the origin. We calculated the potential as

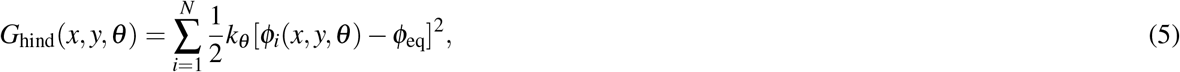

where *G*_hind_ is the harmonic potential for the stator-side angle of the S-F-R connections, *k*_*θ*_ = (64*/π*^2^)*k*_B_*T* is the angular spring constant. Here, *k*_B_ is the Boltzmann constant, *T* is the absolute temperature, and *ϕ*_eq_ = 45° except for Fig. 2e is the equilibrium angle. The stiffness was determined so that the potential of each S is 2*k*_B_*T* at *ϕ*_*i*_ = *ϕ*_eq_ ± 45°.

The equilibrium distribution *P*(*x, y, θ*) is given by the Boltzmann distribution

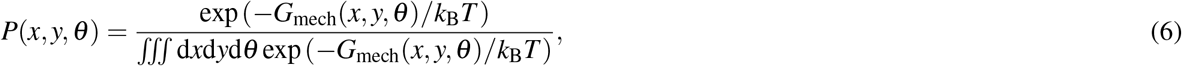

where *G*_mech_ = *G*_ele_ + *G*_elast_ + *G*_hind_ is the interaction potential between the rotor and stator. The trajectories in Fig. 2b–d are the expected values obtained based on *P*(*x, y, θ*).

In Fig. 4, we applied force to evaluate the stall force. We added the free energy *G*_*f*_ (*r, φ*) = *f rφ*. Here, *f* is the magnitude of the hindering force, *r* and *ϕ* are the polar coordinates of the motor.

### Reaction rates

The reactions we consider include the binding and dissociation of DNA strands and the fuel cleavage by the enzyme.

The time scale of all the dynamics is scaled with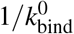, where 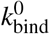is the binding rate of a free F strand to a tethered DNA strand within the volume of (10 nm)^3^. That is, 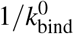 defines the unit time. We estimated 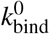to be 500 s^−1^ based on a previous experiment^33^.

We assume that the binding between the tethered strands, R and F-S or F-R and S, occurs only when they are within a cut-off distance *r*_cutoff_ = 12 nm, which is defined as 1.5*L*_eq_. We define a binding rate *k*_bind_(*x, y, θ*) as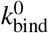 if *r* _*j i*_ (*x, y, θ*) ≤ *r*_cutoff_ and 0 otherwise. Here, *r*_ij_ is the distance between the *i*-th R and *j*-th S.

For the Gillespie algorithm, we used the expected value ⟨*k*_bind_ ⟩ = ∭ d*x*d*y*d*θ P*(*x, y, θ*)*k*_bind_(*x, y, θ*). The dissociation rate is obtained based on the local detailed balance condition. The free energy change associated with the binding events is the mechanical free-energy change *G*_mech_ and chemical free-energy change *G*_chem_. *G*_chem_ is the summation of the free energy change by the hybridization; − *k*_B_*T* for S and F or cleaved product of F, 2*k*_B_*T* for R and F, 6*k*_B_*T* for R and cleaved product of F, which are roughly determined by the typical DNA-binding parameters for the present fuel concentration. The local detailed balance condition yields *k*_dis_(*x, y, θ*) = *k*_bind_(*x, y, θ*) exp[(*G*_mech_ + *G*_chem_)*/k*_B_*T*].

We used the expected value ⟨*k*_dis_⟩ for the Gillespie algorithm.

The binding rate *k*_on_ of free F in the solution to R or S was determined based on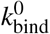. We assume that the binding of a free F strand in the volume of (10 nm)^3^, which corresponds to the concentration of 1.5 mM, is equal to 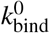. This volume roughly approximates the volume within the cut-off distance *r*_cutoff_ = 12 nm subtracted by the excluded volume due to the rotor and stator structures. We let the concentration of F in the solution to be 3 μM and obtain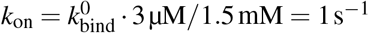. We ignored the rebinding of cleaved products of F to R and S because their concentrations are negligible compared with F. Hybridization is diffusion-limited under the present dilute conditions, and the *k*_on_ does not significantly depend on the sequence^34^. Therefore, *k*_on_ is assumed to be the same for all the sequences. The local detailed balance condition yields sequence-dependent dissociation rate *k*_off_ = *k*_on_ exp(Δ*G*_chem_*/k*_B_*T*).

### Evaluation of run time

When *N* is large, the rotor does not always dissociate within the simulation time. When we observed dissociation in all runs under these conditions, we simply used the mean run time. Otherwise, we estimated the mean run time *T* as follows. Let *Q*(*τ*) be the fraction of the runs in which the motor dissociates from the stator within a time *τ*. Assuming that the time until the dissociation obeys an exponential function *q*(*t*) = *k*_D_*e*^−*t/T*^, we obtain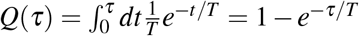. Hence, by evaluating *Q*(*τ*) at a certain *τ*, we obtain *T* = −*τ/* ln(1 − *Q*(*τ*)). We used the longest run time of the runs in which the dissociation takes place within 2 × 10^7^ Gillespie steps as *τ*.

## Acknowledgements

This work was supported by KAKENHI (18H05427, 19H01857, and 23H01136) and JST ERATO (JPMJER2302), Japan.

## Author contributions

K.I. conducted the simulations and analyzed the results, K.I., Y.S., S. T. designed the research and wrote the manuscript. All authors reviewed the manuscript.

## Additional information

The authors declare no conflict of interest.

## Notes

### Competing Interest Statement

The authors have declared no competing interest.

### Summary of Updates

Texts are figures were revised for better readability and correcting errors.

